# Unbiased lineage tracing across the adipose organ decouples developmental origin from anatomical and transcriptomic identities

**DOI:** 10.64898/2026.11.05.762308

**Authors:** Oto Inoue, Archana Singh, Meghana Ginugu, Christopher Ahn, Marine A Secchi, Sarah J Bowling, Hee-Woong Lim, Joan Sanchez-Gurmaches

## Abstract

The developmental origins of individual adipose depots and whether embryonic lineage dictates adult function remain unresolved. Here, we apply *in utero* CRISPR-mediated barcoding lineage tracing to construct the first unbiased clonal map of adipose organ development in mice. The developmental hierarchy we uncover challenges several conventions. Remarkably, interscapular brown fat, used as the prototypical brown fat, is the most developmentally isolated brown fat; depots sharing Myf5/Pax3 lineage markers fail to cluster together; mesenteric white fat shares unexpected high clonal ancestry with brown fats despite a radically divergent transcriptome; and subcutaneous white adipose tissues converge transcriptomically despite distant developmental origins. These findings call for a reinterpretation of single-marker lineage tracing data across the adipose organ and propose that adipose depot identity is not solely encoded by embryonic lineage but is substantially shaped by the anatomical niche.

**Highlights:**

- CRISPR barcoding simultaneously resolves lineage relationships among 12 fat depots.
- iBAT is the most developmentally isolated BAT.
- Visceral white fats are developmentally closer than the subcutaneous depots.
- Developmental origin and transcriptomic identity are largely decoupled across fat depots.

## Introduction

The adipose organ is a multi-depot tissue distributed throughout the body composed of functionally distinct type of adipocytes that individually and cooperatively play indispensable roles in metabolic homeostasis^1,2^. Despite the biomedical significance of depot-specific adipocyte biology, the developmental mechanisms that establish and diversify individual adipose depots remain poorly understood^3,4^.

Adipose tissues are broadly classified into two categories. Brown adipose tissue (BAT) is composed of thermogenic brown adipocytes rich in mitochondria that express uncoupling protein 1 (UCP1). White adipose tissue (WAT) is formed by white adipocytes with few mitochondria, organized into subcutaneous and visceral depots^5^. A third adipocyte type, the thermogenic beige or brite cell, also express UCP1 and can be induced mainly within subcutaneous WAT depots under specific stimuli^6,7^. This is thought to, at least partially, contribute to the distinct intrinsic properties of subcutaneous and visceral WAT depots and their distinct contribution to metabolic disease^8–12^. Although this classification provides a useful framework, it belies the remarkable functional specialization emerging at the level of individual depots: each adipose depot exhibits a distinct transcriptional, metabolic, and secretory profile closely matched to the physiological demands of its anatomical niche. Although these classifications help describe key features of adipocytes, they do not address the ontogenetic relationships among adipocytes or depots.

All adipocytes were assumed to originate from a common mesodermal lineage, a view that was inherited from the classical histological designation of adipose tissue as a connective and inert tissue^4,13–15^. However, over the past two decades, Cre/Lox-based lineage tracing in mice has revealed an unexpectedly complex set of developmental relationships both between and within individual adipose depots. First, the same adipocyte types can originate from distinct progenitor pools (eg, interscapular BAT adipocytes trace to a Myf5^+^ lineage while adipocytes in periaortic BAT do not^16–19^). Second, individual depots can be generated from more than one progenitor lineage (eg, half of adipocytes in male perigonadal WAT originate from a somitic mesoderm population traced by Meox1-Cre expression^20^). Third, distinct types of adipocytes can share developmental markers (eg, Myf5-Cre and Meox1-Cre label subsets of both brown and white adipocytes^16,20–23^). Fourth, the adipose organ is not exclusively generated from a solely specialized area of the mesoderm (eg, some cephalic fat depots and periaortic BAT receive substantial contributions from neural crest progenitors of ectodermal origin^17,24^). Collectively, these findings establish that adipocyte identity is not determined by a single progenitor lineage but emerges from the convergence of multiple, germ layer–spanning developmental programs whose intersection may be unique to each depot.

Despite these advances, fundamental questions regarding the development of the full complement of adipose tissue depots remain unresolved in part due to intrinsic limitations of the systems used. Cre/lox-based fate mapping and conventional binary reporters trace a single lineage per experiment. This makes it impossible to determine whether adipocytes sharing a particular developmental marker descend from a common progenitor or from independently progenitor populations that co-express that marker. Likewise, the relationship between adipocytes not originating from a particular developmental marker cannot be determined. Thus, the limited resolution of the systems used left open the fundamental question of the ontogenetic history and relationship between each fat depot.

Here, we leverage a CRISPR-mediated barcoding reporter mouse model (CARLIN)^25^ to perform the first comprehensive, unbiased clonal analysis of adipose tissue ontogeny across twelve major BAT and WAT depots. By activating barcoding during embryonic development and analyzing clonal representation across depots at maturity, we construct a unified lineage map that reveals shared and unique clonal origins and identifies previously unrecognized ontogenetic relationships between adipose tissues. We coupled this developmental analysis with functional status at the morphologic and transcriptomic levels. In summary, our findings provide a high-resolution developmental framework and its relationship to functional outcomes of the whole adipose organ.

## Results

### Morphological analysis of adipose depots reveals signatures that are discordant with known developmental origins

Morphological differences among adipocytes underlay functional modifications. To characterize the morphological heterogeneity of discrete adipose tissue depots, we performed quantitative histological analysis on eight-week-old C57BL/6J male mice. We quantified nuclear density (nuclei/mm²) across six brown adipose tissue (BAT) depots: interscapular (iBAT), subscapular (sBAT), cervical (cBAT), aortic arch (aaBAT), thoracic aorta (taBAT), and perirenal (prBAT) [Figure 1A]. In parallel, we measured adipocyte cross-sectional diameter (µm) across six white adipose tissue (WAT) depots: posterior subcutaneous (psWAT), mesenteric (mWAT), perigonadal (pgWAT), retroperitoneal (rWAT), dorsal anterior subcutaneous (dasWAT), and ventral anterior subcutaneous (vasWAT) [Figure 1A].

**Figure 1:**
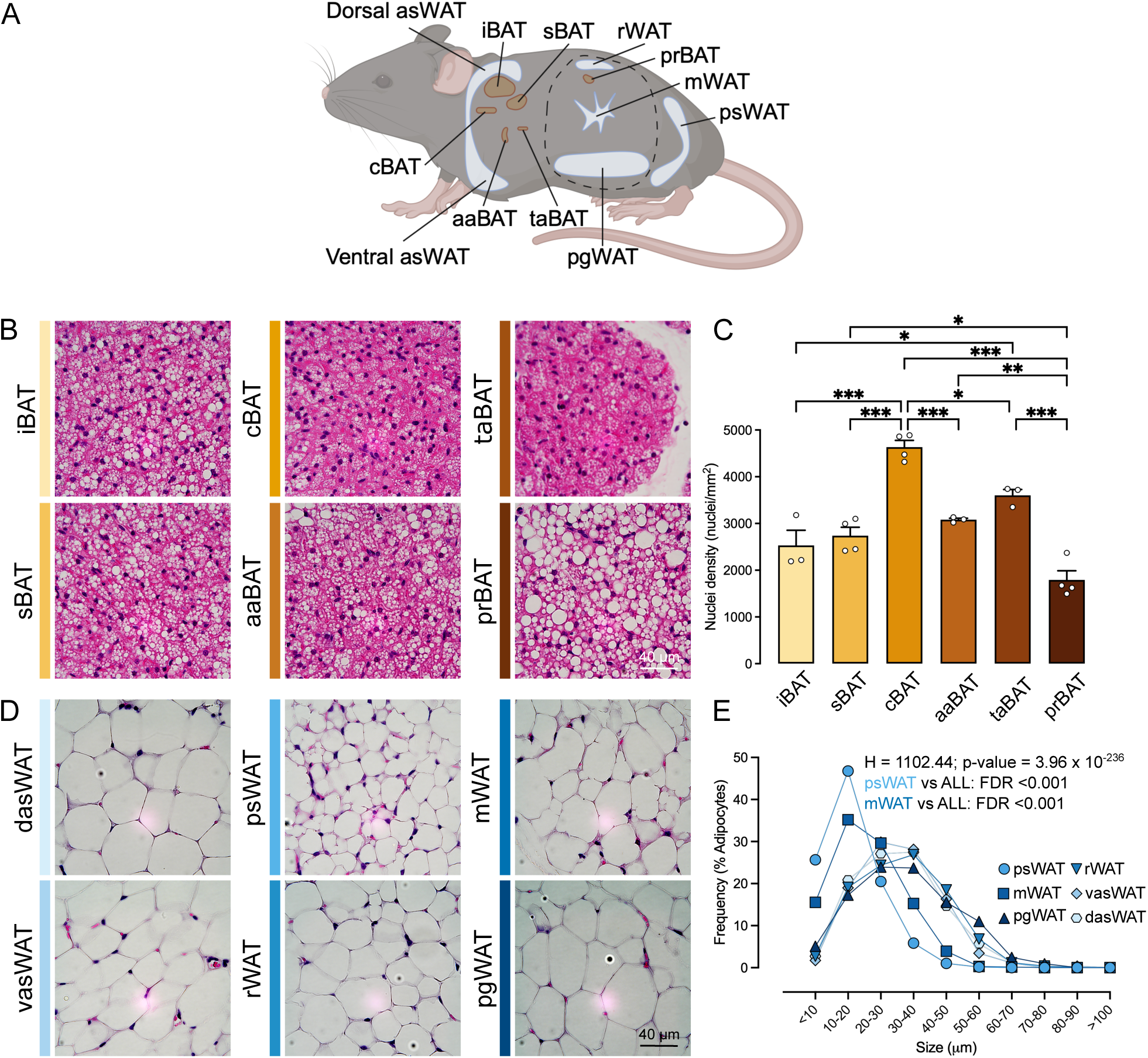
Morphological heterogeneity across BAT and WAT depots does not recapitulate known developmental relationships. (A) Diagram of the adipose organ depicting the depots analyzed in this study. CreCreated in BioRender. Sanchez Gurmaches, J. (2026) https://BioRender.com/bcak2da. (B) Representative H&E images of BAT depots of 8-weeks-old C57Bl/6J male. n= 3-4 per depot. (C) Nuclei density quantification from H&E images of BAT depots of 8-weeks-old C57bl/6J male. n= 3-4 per depot. Data are mean + SEM, one-way ANOVA followed by Tukey’s multiple comparisons test. *P < 0.05, **P < 0.01, ***P < 0.001. (D) Representative H&E images of WAT depots of 8-weeks-old C57Bl/6J male. n= 4 per depot. (E) Adipocyte size distribution in WAT depots of 8-weeks-old C57Bl/6J male. n= 4 per depot. Kruskal-Wallis test followed by Dunn’s test with Benjamini-Hochberg FDR correction.

Nuclear density differed significantly among BAT depots [Figure 1B-C]. cBAT exhibited the highest nuclear density, significantly exceeding all other depots except aaBAT, which may reflect unique structural or functional properties of these two depots [Figure 1B-C]. In contrast, prBAT displayed the lowest nuclear density and contained visibly larger lipid droplets. iBAT, sBAT, aaBAT, and taBAT occupied intermediate positions [Figure 1B-C]. Notably, this morphological hierarchy does not recapitulate some of the established developmental relationships among these depots.

To assess morphological heterogeneity among WATs, we quantified adipocyte cross-sectional diameter [Figure 1D-E]. Comparison among all datasets revealed significant differences among depots (H = 1,102.44, p = 3.96 × 10^-2^^36^), and pairwise comparisons identified three statistically distinct tiers [Figure 1D-E]. psWAT contained the smallest adipocytes, consistent with the established browning propensity and higher thermogenic capacity of this depot. mWAT exhibited intermediate adipocyte size, significantly larger than psWAT and significantly smaller than all remaining depots. pgWAT, rWAT, dasWAT, and vasWAT formed a homogeneous group harboring the largest adipocytes [Figure 1D-E]. This morphological convergence is striking given the known differences in anatomical location, function and developmental origin among these four depots.

Together, these morphometric data demonstrate that adipose organ depots exhibit profound, depot-specific morphological signatures. Critically, these signatures do not align with the developmental relationships established by lineage-tracing studies, suggesting either that unknown developmental relationships exist or that postnatal functional adaptation and local microenvironmental cues, rather than embryonic origin alone, may be determinants of adipocyte morphology.

### Transcriptomic identity does not track known developmental origin across the adipose organ

The pronounced morphological differences among similarly classified depots suggested underlying functional divergence. To test this using transcriptomic identity as a proxy for function, we profiled all twelve depots from eight-week-old C57BL/6J males by bulk RNA-seq. Principal component analysis cleanly separates BAT from WAT, consistent with the fundamental divergence between these classes [Figure S1A]. The two largest upregulated modules across BAT over WAT are dominated by mitochondrial bioenergetic programs (eg, thermogenesis, oxidative phosphorylation, electron transport chain) [Figure S1B]. Conversely, the largest WAT-enriched module over BAT is dominated by a large variety of programs (eg, PI3K/Akt signaling, extracellular matrix and macrophage chemotaxis) [Figure S1B].

To resolve functional heterogeneity within BAT at a higher resolution, we analyzed the six BATs independently. PCA and Spearman correlation consistently group them into three clusters [Figure 2A-B]. iBAT, sBAT, and cBAT form a compact triplet with the highest pairwise correlations in the dataset (ρ = 0.989–0.993) in line with their common dermomyotome proposed origin^16,19,21^; taBAT and aaBAT pair together (ρ = 0.970) aligning with their common neural crest origin^17^; and prBAT segregates as the most divergent BAT depot (ρ = 0.904–0.925) despite a putative origin in the dermomyotome showed by shared embryonic expression of *Pax3* to iBAT, sBAT and cBAT^16^. Notably, the canonical thermogenic genes (ie, *Ucp1*, *Prdm16*, *Cidea*, *Ppargc1a*, *Dio2*, and *Elovl3*) are not differentially expressed among the depots, indicating that they define a shared BAT identity rather than inter-depot variation.

**Figure 2:**
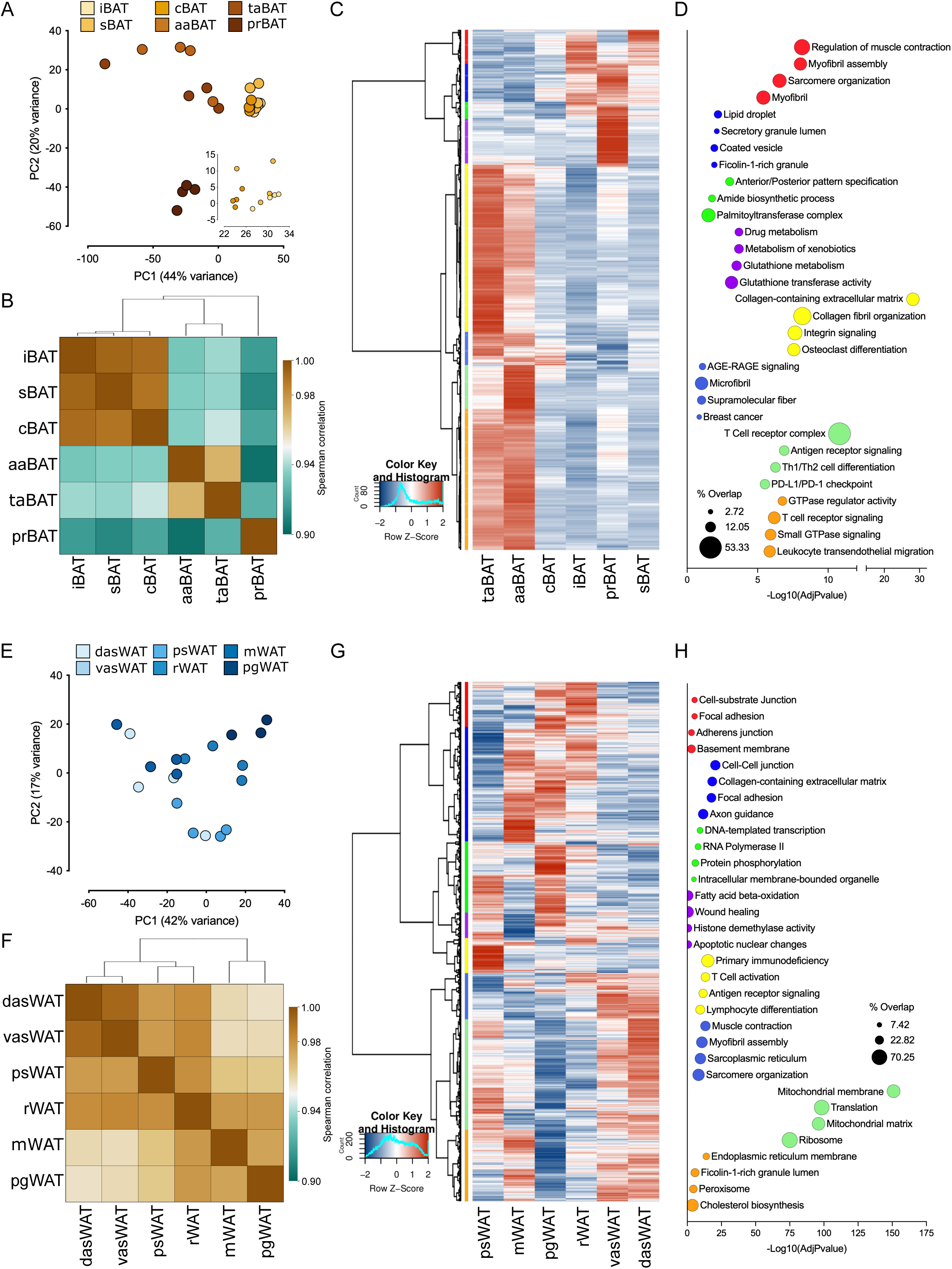
Transcriptomic profiling reveals depot-specific gene expression programs with strong influence of functional axes independently of adipocyte type. (A) PCA analysis of BAT depots of 8-weeks-old C57Bl/6J male. Each point represents one biological replicate, colored by depot. n= 4 per depot. (B) Spearman correlation matrix and clustering across average values of BAT depots of 8-weeks-old C57Bl/6J male. n= 4 per depot. (C) Hierarchical clustering of averaged depot transcriptomes of BAT depots of 8-weeks-old C57Bl/6J male. Co-expression modules by unsupervised clustering are indicated by the colored bar at left. n= 4 per depot. (D) Gene Ontology and pathway enrichment by module of BAT depots of 8-weeks-old C57Bl/6J male. Dot color matches cluster colors in panel C. (E) PCA analysis of WAT depots of 8-weeks-old C57Bl/6J male. Each point represents one biological replicate, colored by depot. n= 4 per depot, except pgWAT n=3. (F) Spearman correlation matrix and clustering across average values of WAT depots of 8-weeks-old C57Bl/6J male. n= 4 per depot, except pgWAT n=3. (G) Hierarchical clustering of averaged depot transcriptomes of WAT depots of 8-weeks-old C57Bl/6J male. Co-expression modules by unsupervised clustering are indicated by the colored bar at left. n= 4 per depot, except pgWAT n=3. (H) Gene Ontology and pathway enrichment by module of WAT depots of 8-weeks-old C57Bl/6J male. Dot color matches cluster colors in panel G.

Next, we analyzed gene modules attributed to particular BAT depots by unbiased clustering. The two largest modules, yellow (645 genes) and orange (541 genes), together exceed 60% of all differentially expressed genes and are elevated in taBAT and aaBAT [Figure 2C-D, S1C]. The yellow module, highest in taBAT, is enriched for immune, inflammatory, and extracellular matrix pathways, and the orange module, highest in aaBAT, reinforces this immune axis with adaptive immune response and lymphocyte differentiation. A third taBAT/aaBAT module (light green, 168 genes) contains the myokine Fndc5/irisin and is enriched for calcium signaling, and a fourth (royal blue, 127 genes) for fibroblast growth factor receptor signaling. Together these modules reveal that taBAT and aaBAT harbor a markedly more complex immune and stromal microenvironment than the other BATs, likely reflecting a perivascular identity associated to their anatomical proximity to the aorta. In contrast, the module defining the iBAT/sBAT/cBAT triplet (red, 131 genes) [Figure 2C-D], highest in iBAT and sBAT [Figure S1C], retains a skeletal muscle contractile and de novo lipogenesis signature. This co-occurrence underscores the proposed shared ontogeny between classical brown adipocytes and the dermomyotomal lineage. Finally, three modules selectively upregulated in prBAT (purple, green, blue) are dominated by developmental patterning, including anterior-posterior specification and hedgehog signaling, alongside lipid metabolic programs such as lipoprotein metabolism, cholesterol metabolism, and long-chain fatty acid transport [Figure 2C-D, S1C]. Adipose tissues express patterning genes depending on their anatomical position maybe independently of their functions^26^. However, this convergence of developmental and lipid programs sets prBAT apart from all other brown depots.

Applying the same approach to the six WAT depots, PCA separates subcutaneous (psWAT, dasWAT, vasWAT) from visceral (pgWAT, mWAT, rWAT) depots, with rWAT at an intermediate position [Figure 2E]. Spearman correlation confirms that dasWAT and vasWAT are near-identical (ρ = 0.992) despite originating from independent lineages (ie, Myf5-Cre and Prx1-Cre, respectively)^16,20,276,20,27^, pgWAT and mWAT anchor the visceral pole (ρ = 0.974) even all lineages associated to pgWAT adipocytes are not shared by mWAT^16,20,276,20,27^, and rWAT bridges the two compartments albeit it shares a common Myf5-Cre and Pax3-cre lineage with dasWAT^16,20,23,27^ [Figure 2F]. The lowest WAT coefficients, between pgWAT and dasWAT and vasWAT (ρ = 0.956–0.958), still exceed 0.95 [Figure 2F]. Thus, inter-depot divergence within WAT is substantially narrower than within BAT (minimum ρ = 0.904), indicating that BATs are the more transcriptionally specialized class [Figure 2B, 2F].

Unsupervised clustering resolves eight WAT modules [Figure 2G-H, S1D]. The largest (light green) is upregulated in subcutaneous relative to visceral depots and dominated by mitochondrial bioenergetic programs, including cellular respiration, oxidative phosphorylation, and thermogenesis, containing electron transport subunits, thermogenic effectors (ie, *Ucp1*, *Ppargc1a*, *Prdm16*), and fatty acid oxidation machinery (ie, *Cpt1b*, *Cpt2*, *Acadvl*). This is consistent with the known browning capacity of subcutaneous WATs and the low thermal stress experienced by mice housed at 22°C. A second module (orange, 1,200 genes), highest in mWAT, dasWAT, and vasWAT and suppressed in pgWAT, is enriched for cholesterol biosynthesis and fatty acid metabolism. The royal blue module, highest in vasWAT and dasWAT, is enriched for muscle pathways such as muscle contraction and sarcomere organization. dasWAT has a shared Myf5/Pax3 lineage origin with muscle which may explain this result, but that is not true for vasWAT^16,19,23^. The blue module, highest in mWAT and pgWAT, is enriched for cell migration, cell-matrix adhesion, and heart development, and the red module, shared by rWAT and pgWAT, for extracellular matrix and angiogenesis. Strikingly, the immune-enriched yellow module is most strongly expressed in psWAT, far exceeding all other depots, likely reflecting its proximity to the inguinal lymph node (which was removed from psWAT before analysis) and associated lymphatic drainage rather than the metabolic inflammation typical of visceral fat. The green module, enriched for Hox transcription factors, is highest in pgWAT and psWAT, and the purple module, enriched for fatty acid beta-oxidation and lipolysis (*Lipe*, *Plin1*), is upregulated in pgWAT and psWAT but profoundly suppressed in mWAT, indicating depot-specific lipolytic capacity. Together, these data show that transcriptomic divergence among WAT depots is driven by mitochondrial and lipogenic metabolism, extracellular matrix and immune composition, and developmental programs.

Taken all together, we observe a strong mismatch between transcriptomic identity and known developmental relationships.

### Unbiased lineage tracing uncovers unexpected developmental relationships among fat depots

The poor alignment between transcriptomic and morphological data with known developmental origins we showed above prompted us to revisit the lineage relationships among the adipose organ, moving beyond the limited resolution of traditional single-marker tracing studies.

To map the developmental relationships among adipose depots in an unbiased manner without relying on candidate markers, we performed *in utero* CARLIN lineage barcoding^25^. To use this model, we crossed double homozygous transgenic males with globally expressed Tet-inducible Cas9 and a reverse tetracycline controlled transactivator with females homozygous for the globally expressed CARLIN transgene [Figure 3A]. In the CARLIN model, CRISPR-Cas9 uses predesigned 10 gRNA towards a target within the transgene that is formed by 10 repetitions in tandem of cutsides within the same transgene but not specific to any other areas of the mouse genome [Figure 3B]. This generates semi-random barcodes within the CARLIN transgene that are transcribed thanks to the strong CAG promoter and can be detected by RNA-seq upon making libraries that are specific to the target sequence within the CARLIN transgene. The inheritance of these barcodes allows the developmental relatedness of tissues to be inferred from the barcodes that their constituent cells share. We induced Cas9 activity with three pulses of doxycycline through retroorbital injections to timed pregnant dams at E6.5, E9.5, and E13.5 to label progenitors as they colonize and give rise to the distinct adipose depots [Figure 3A]. First, we assessed editing efficiency in the CARLIN males subjected to our doxycycline protocol. We found that doxycycline induced robust editing, with 50% of sequences modified in fat and 25-50% of modifications in other tissues [Figure S2A], similar to previously determined editing rates^25^. Next, we isolated RNA from the above twelve adipose tissues of three independently labelled eight-week-old male mice and generated CARLIN array libraries. Barcodes shared between mice arose independently and our filtering strategy reduced this overlap [Figure S2B]. The majority of barcodes recovered from every tissue were significantly enriched relative to an allele bank with all previously generated alleles^25^, as expected [Figure S2C-D]. Because each animal acquires a distinct barcode repertoire, we analyzed every mouse independently, generating a consensus tree from 10,000 stochastic reconstructions to capture the hierarchy of clonal relatedness across depots in each mouse [Figure S2E-G]. We then integrated these results into a pairwise tissue distance matrix that provides a quantitative summary of the developmental relationships among adipose depots.

**Figure 3:**
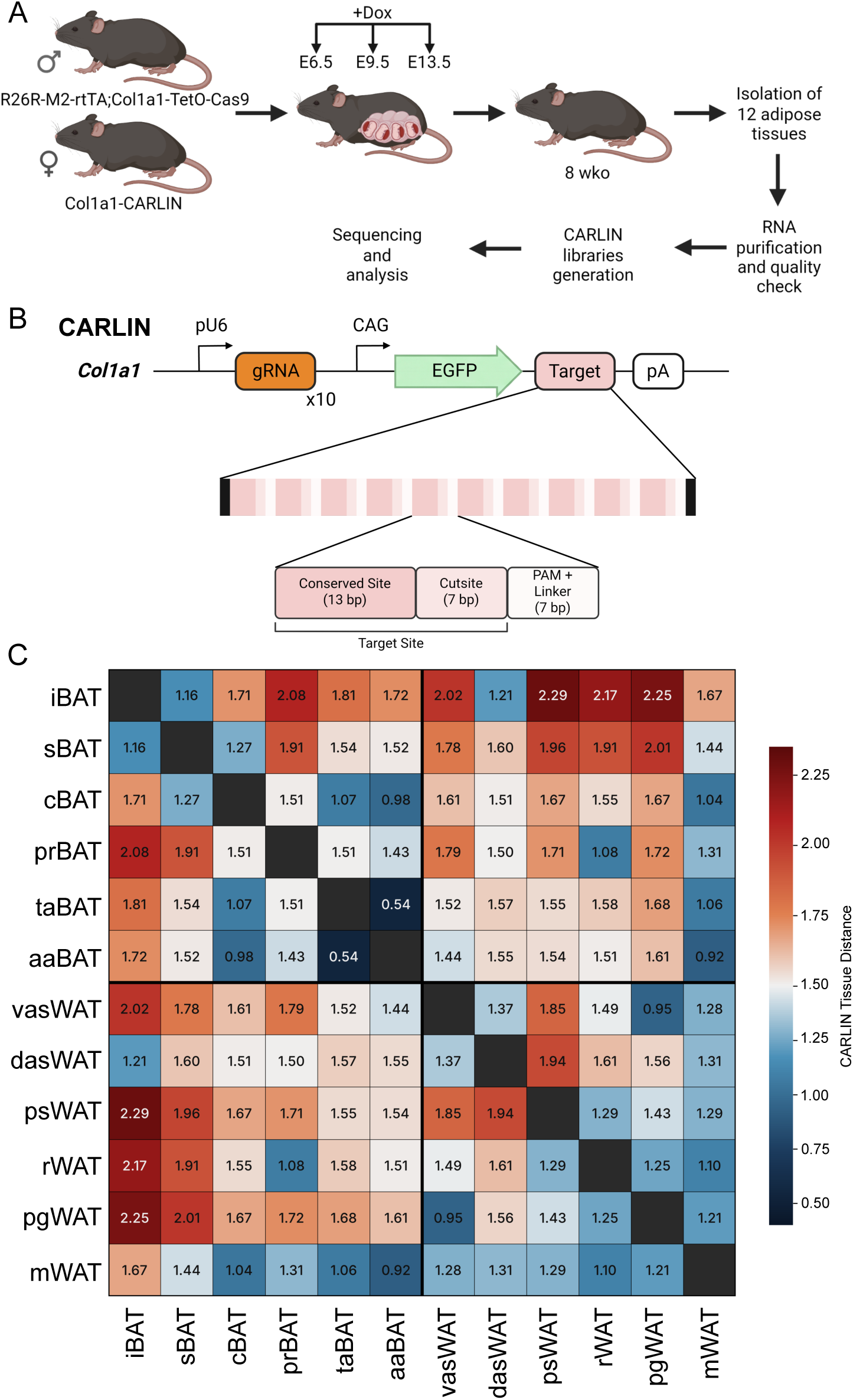
CARLIN barcoding uncovers an unexpected developmental hierarchy across the adipose organ. (A) Diagram of the breeding strategy and experimental plan for CARLIN mice experiments. Created in BioRender. Sanchez Gurmaches, J. (2026) https://BioRender.com/kt0ftxh. (B) Diagram of the CARLIN transgene. The target of the 10 gRNA generated is divided in an array of 10 motifs each divided into a 13-bp conserved site (out of range of Cas9 editing), a 7-bp cutsite and a protospacer adjacent motif (PAM) and linker final motif. Created in BioRender. Sanchez Gurmaches, J. (2026) https://BioRender.com/nebo30f. (C) CARLIN tissue distance matrix among twelve adipose tissues. n=3.

This analysis reveals an organization that recapitulates several established relationships while uncovering unexpected ones [Figure 3B]. Among the BAT depots, the smallest developmental distance separates aaBAT and taBAT, which form the closest pair in the entire dataset. The next closest relationships are those between aaBAT/cBAT and taBAT/cBAT, both of which are closer than the iBAT/sBAT pair. This is unexpected, because iBAT, sBAT, and cBAT are conventionally grouped as true classical BAT^28,29^. In our data, however, the distances from iBAT and sBAT to cBAT are considerably larger than this prevailing grouping would predict. This is partially consistent with iBAT, sBAT and cBAT completely sharing developmental Pax3-Cre expression but cBAT only partially sharing Myf5-Cre lineage with iBAT and sBAT^16^. The two greatest distances between any BAT depots both involve prBAT, in its comparisons with iBAT and sBAT, yet prBAT shows only mild and notably uniform distances to cBAT, aaBAT, and taBAT. Most strikingly, iBAT emerges as the single most developmentally isolated depot in the dataset, showing higher developmental distance from every other BAT depot, with the sole exception of sBAT, than any other BAT. Given that iBAT is by far the most studied BAT, this developmental isolation argues for caution in treating it as representative of BAT.

Among the WAT depots, the visceral depots (rWAT, pgWAT, and mWAT) are on average developmentally closer to one another than are the subcutaneous depots (dasWAT, vasWAT, and psWAT) [Figure 3B]. Notably, mWAT shows a small developmental distance to every other WAT depot, irrespective of visceral or subcutaneous anatomical location. The smallest distance among the WAT depots is that between pgWAT and dasWAT, a pairing of one visceral and one subcutaneous depot. This is surprising but partially recreates the established partially shared Pax3-Cre lineage with among these two fat depots^16,22^. Unexpectedly, the distance separating the two asWAT exceeds any distance measured between visceral depots. The largest distances among the WATs are those between psWAT and dasWAT, followed by psWAT and vasWAT. Unexpectedly, the greatest distance for rWAT is to dasWAT, even though the two share a Myf5-derived developmental origin^16,21^.

The developmental distances between WAT and BAT depots offer further insight into the broader architecture of adipose tissue development, a relationship that earlier studies had already shown to be unexpectedly complex. mWAT has the lowest average developmental distance to the BAT depots of any WAT analyzed, despite the absence of any prior report of shared lineage [Figure 3B]. Conversely, rWAT shows some of the greatest distances in the entire dataset in its comparisons with iBAT and sBAT, even though earlier work indicated that these depots share a dermomyotomal origin marked by Myf5 expression. The smallest distance between dasWAT, another depot reported to arise from the Myf5 lineage, and any BAT depot is to iBAT. rWAT, in turn, displays substantial clonal overlap with prBAT, pointing to a shared developmental niche even though the two depots are populated by distinct adipocyte types. Finally, pgWAT exhibits the largest average developmental distance from the BAT depots while remaining generally close to the other white depots.

Taken together, this unbiased reconstruction of the developmental relationships among murine adipose depots reveals organizational features that challenge assumptions drawn from earlier, marker-restricted lineage-tracing studies and indicates that anatomical classification alone is an incomplete predictor of developmental relatedness.

### Developmental lineage and transcriptomic identity are substantially decoupled

A long-standing assumption in adipose biology is that depot function is dictated by developmental origin. This is best exemplified by the paradigm that the high metabolic rate of BAT stems from its shared lineage with muscle, even though not all brown adipocytes follow this trajectory^16,19,30^. Leveraging our unbiased developmental map, we asked to what extent the transcriptomic identity of a depot, as a proxy for its function, is set by lineage rather than by other factors (eg, postnatal microenvironment).

To do this, we generated a Spearman rank correlation matrix from the developmental and gene expression distance matrices [Figure 2-3]. This revealed a weak correlation between developmental and transcriptomic similarity (r = 0.345, p = 0.004). Lineage therefore accounts for a minor fraction of transcriptomic variation, indicating that the two are substantially decoupled. To resolve this decoupling pair by pair, we integrated the gene expression and CARLIN barcoding distances for every pair of depots into a single discordance matrix [Figure 4]. In this analysis, positive values denote depots more similar by expression than by lineage (transcriptomic convergence) and negative values denote the reverse (transcriptomic divergence).

**Figure 4:**
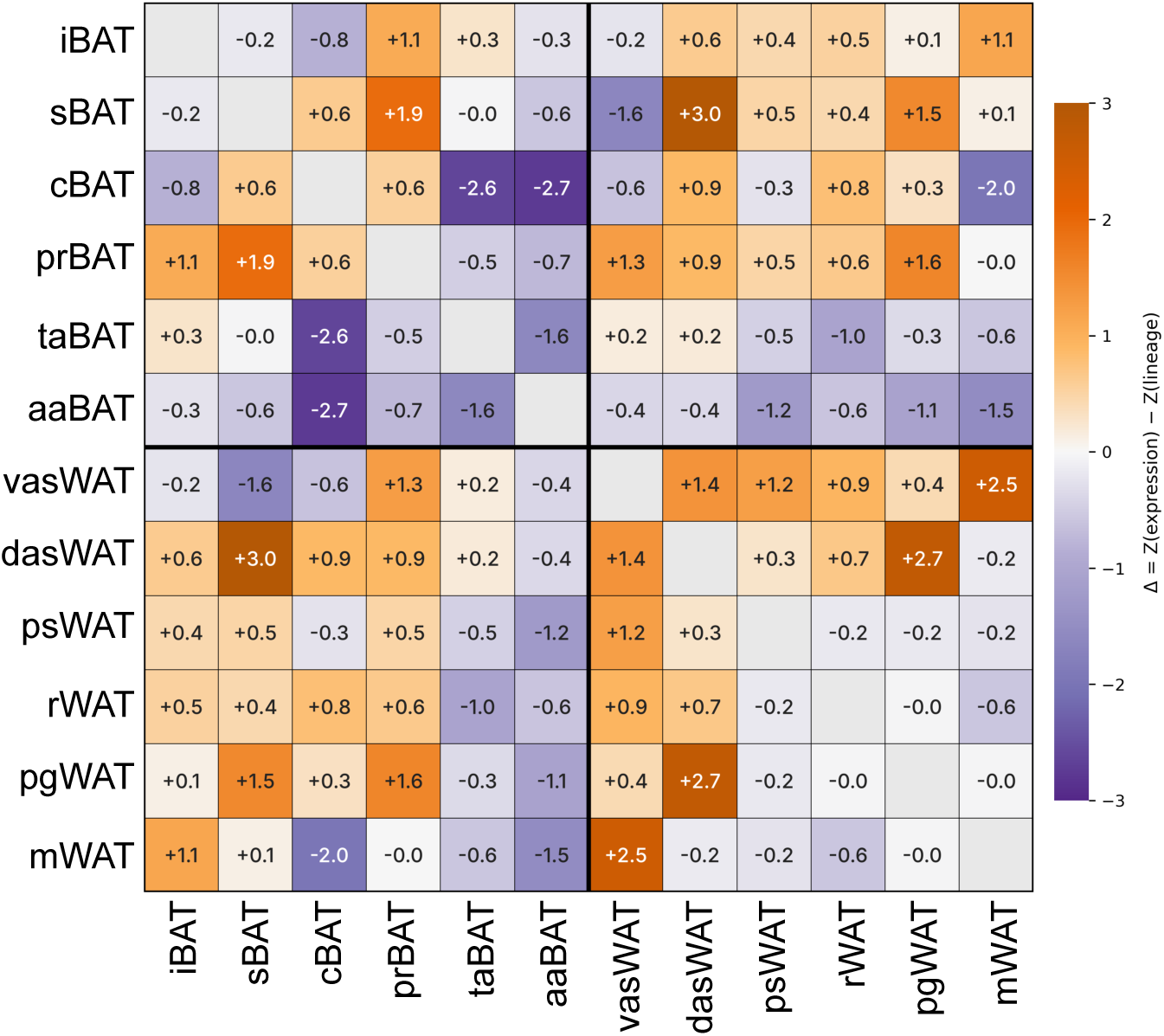
Discordance between transcriptomic similarity and developmental relatedness is depot dependent. Discordance matrix between gene expression and development of adipose tissues of 8-weeks-old mice. n=3.

In some pairs, the two measures align. The low discordance between iBAT and sBAT reflects both their shared developmental origin, confirmed here without bias, and their closely matched transcriptomes. At the other extreme, sBAT and vasWAT differ substantially yet proportionately in both dimensions.

The strongest convergence occurs among subcutaneous WAT depots: dasWAT and psWAT show the highest discordance index (Δ = +2.98), followed by vasWAT and psWAT (Δ = +2.74), pairs with close transcriptomes despite distant developmental origins. cBAT and iBAT likewise converge transcriptionally (Δ = +2.53) despite being developmentally distant. Notably, dasWAT and rWAT, both reported to derive from a Myf5/Pax3 lineage^16,23^, converge transcriptionally beyond their developmental distance (Δ = +1.93). This illustrates that a shared single-marker lineage does not predict either unbiased developmental relatedness or transcriptional coordination.

The most striking divergences involve mWAT and the brown depots cBAT, taBAT, and aaBAT (Δ = −1.96, −2.62, and −2.75), which share substantial clonal ancestry yet adopt radically different adult transcriptomes. prBAT and rWAT serve as an internal control: their shared perirenal developmental niche is matched by a low CARLIN distance (1.078) and elevated transcriptomic similarity (ρ = 0.952), yielding a near-concordant (Δ = +0.49). This identifies the perirenal fat pad as a genuinely intermediate compartment where lineage and expression remain coupled.

Together, these results indicate that lineage-independent mechanisms can override the developmental origin of a depot to shape its mature identity in a niche dependent manner.

## Discussion

The developmental architecture of the adipose organ remains incompletely resolved^3,4^. This is perhaps most evident in our inability to fully explain the developmental divergence between even the most distinct adipocyte classes, such as classical brown and white adipocytes^16,19,23^. In addition, the developmental logic that assigns functionality to each adipose depot remains poorly understood.

The broadly established single-marker tracing and binary reporter systems (eg, Cre/reporter systems) carry increasingly appreciated limitations in developmental biology. Critically, these designs interrogate a single lineage per experiment, cannot distinguish whether adipocytes that share a developmental marker descend from a common progenitor or from independent progenitor pools that co-express a marker, and they are equally uninformative about the relationships among adipocytes that lack the marker. Cre-mediated lineage tracing carries additional, well-documented, liabilities. Different cells may express Cre at different levels, there may be off-target expression from the transgene or its flanking sequences, and the genomic insertion site of most drivers is unknown^31–41^. Inducible drivers add further uncertainty, because CreER lines exhibit tamoxifen-independent basal recombination^42^ and because tamoxifen persists at active concentrations for weeks, eroding the temporal precision of pulse labeling^43,44^. Together, these constraints have left the ontogenetic architecture of the adipose organ only partially mapped.

The CARLIN system, applied here to adipose development, circumvents many of these constraints^25^. First, it is unbiased, requiring no preselection of markers. Second, it bypasses the problems associated with using multiple Cre-drivers. Third, it resolves the developmental relationships among each tissue/cell analyzed within a single animal, a comparison that no Cre/loxP experiment can deliver.

Among the most recognized frameworks in adipose tissue development is that subsets of BAT and WAT share a common Myf5/Pax3 lineage^16–19^. This includes all adipocytes in the iBAT, sBAT, rWAT, and dasWAT. However, these depots do not cluster together by CARLIN distance, prompting a reassessment of this model. rWAT in particular lies among the most developmentally distant depots from iBAT and sBAT in the entire dataset. This may indicate that some lineage markers report anatomical region of origin rather than ontogenetic proximity. The cells that expressed Myf5-Cre that generate the iBAT or dasWAT and those that generate the rWAT may share a marker but not necessarily be the same common ancestor, even if originating from within the same developmental region.

iBAT serves as the prototype for BAT in the field, and much of what is known about BAT biology derives from this single depot. Our unbiased map argues against that generalization. We found that iBAT has the greatest developmental distance to every other BAT, except sBAT. This is consistent with our data here and previous discoveries with candidate-marker tracing that indicate that multiple populations of progenitors exist in the dermomyotome and they selectively contribute to distinct BAT depots. For example, iBAT and sBAT, are fully labelled by Myf5-Cre while prBAT is not ^16,19,23^. However, tracing with the additional dermomyotome marker Pax3-Cre labels all brown adipocytes in iBAT and most of them in prBAT^16^. Meox1-Cre, another somitic marker, also traces to iBAT, but its contribution has not been confirmed in other BAT depots, so whether this represents a pan-brown adipocyte marker is to be determined^20^. iBAT also differs substantially in gene expression from BAT in not classically tested locations (ie, aaBAT, taBAT and prBAT). These two factors reinforce that conclusions drawn from iBAT alone may not extend to other, far less studied BAT depots.

The periaortic BAT depots, aaBAT and taBAT form the closest developmental pair in our dataset. Unexpectedly, cBAT sits developmentally closer to aaBAT/taBAT than to iBAT/sBAT. This dissociation is not without precedent. The iBAT and sBAT Myf5-lineage is known to be incompletely shared by cBAT^16,19,23^. aaBAT possesses a distinct progenitor resident cell^45^, and it traces largely to neural crest rather than to paraxial mesoderm^17^. This extends earlier data showing that cephalic white adipocytes surrounding the salivary glands derive from neural crest^24^. This fragments the conventional grouping of iBAT, sBAT, and cBAT as homogeneous classical BAT, a grouping that rests on shared gene expression rather than on shared ancestry.

mWAT shows the lowest average developmental distance to the BAT depots of any white depot even though its adult transcriptome is radically different. This is unexpected due to the previously reported distinct origins of mWAT (splanchnic posterior lateral plate mesoderm (HoxB6^+^)), iBAT/sBAT (dermomyotome) and of aaBAT/taBAT (neural crest)^20^. The transcriptome of mWAT also indicates that despite a close ontogenetic relationship to all fats analyzed, mWAT has a unique function as a primary immunological barrier for the gut. On the other hand, the proposed ontogenetic origin of mWAT with psWAT and female pgWAT is reaffirmed with a low distance score in our dataset (albeit our data are in male mice)^20^. However, these distances are not very different than those seen for other WATs from other compartments. This new information expands our incomplete understanding of the development of mWAT. We hypothesize that mWAT illustrates a model that developmental origin may set the starting point, but the microenvironment has a strong pull in the adult transcriptional and functional identity.

The subcutaneous depots make the complementary case. dasWAT and psWAT show the largest positive discordance in the dataset, followed by vasWAT and psWAT. This is due to near-identical transcriptomes across depots that CARLIN distance places far apart. We interpret that the subcutaneous cluster is an example of the microenvironment imposing a convergent transcriptome on depots of distinct origin. Previous Cre-mediated lineage tracing showed that adipocytes within psWAT and vasWAT shared a common progenitor that expresses *Prx1-Cre*, while those in dasWAT do not^27^. The subcutaneous niche is characterized by dermal proximity, lymphatic drainage, innervation patterns, and mechanical properties that are shared across subcutaneous depots regardless of their anterior-posterior position^46–49^. Our new data suggest this transcriptomic convergence is not a consequence of shared ancestry. Together, the mWAT and subcutaneous WAT cases of distinct directions of convergences constitute an argument for niche dominance in adult adipose tissue transcriptome establishment.

prBAT and rWAT share a high CARLIN proximity and high transcriptomic similarity despite belonging to different adipose classes yielding low discordance. This suggests that fat pads in close proximity may show highly coupled lineage and transcriptome despite being categorized as distinct types of adipose tissue (BAT vs WAT). The perirenal compartment is a clinically translatable depot because in humans the perirenal region harbors widespread dormant brown adipocytes and progenitor cells that retain the capacity to be reactivated^50^. This raises the need to better understand prBAT to exploit it to influence whole-body metabolism.

In this study, we recovered CARLIN barcodes from whole tissue rather than from adipocytes alone, which may affect our data interpretation. Single-nucleus resolution would be preferable but is not currently feasible in this model. The transgene includes a nucleus export signal^25^ that prevents barcode retention in nuclei, and whole-adipocyte single-cell capture at scale remains technically impractical for buoyant mature adipocytes. Our analysis was restricted to male mice. Thus, it cannot address sex differences. Sex differences are well documented for the lineages that build pgWAT in particular^16,20^. Depot-specific sexual dimorphism in adipose development is a natural target for future high-resolution barcoding studies. Ambient temperature deeply affects adipose tissue structure and transcriptomic pattern. Here, we analyzed C57Bl/6J mice raised and maintained at 22°C which is below thermoneutrality. The changes in the relationships between gene expression and development associated to ambient temperature could be an exciting area of future research.

In conclusion, we provide an unbiased view of the ontogenetic relationships across a broad set of adipose depots. This proves the feasibility and possibilities of using new CRISPR-based lineage barcoding systems to further understand adipose tissue biology. In doing so, we revise several assumptions inherited from marker-restricted lineage tracing. The field has assumed that developmental origin largely dictates depot function. Our data indicate instead that origin accounts for only part of functional identity and that much of the remainder might be written by the anatomical niche. This does not render lineage irrelevant. When lineage and niche align, they reinforce one another (eg, prBAT/rWAT pair). When lineage and niche conflict (eg, among subcutaneous depots), the niche prevails in setting the functional outcome. The use of new lineage tracing approaches in adipose tissue introduced here provides a route toward the depot-selective tools that the field has long lacked. Our new framework reshapes how adipose development can be interpreted.

## Methods

### Lead contact and material availability

Further information and request for resources and reagents should be directed to and will be fulfilled by the Lead contact, Joan Sanchez-Gurmaches.

### Mice and mice housing

All mice used in this study were in C57Bl6/J background. CARLIN (B6;129S4-Col1a1tm4(CAG-EGFP)Fcam/Mmjax, JAX stock 067061, backcrossed in house for 10 generations to C57Bl6/J), TetO-Cas9 (Col1a1-tm1(tetO-cas9)Sho, JAX stock 029476), rtTA-M2 (Gt(ROSA)26Sor-tm1(rtTA*M2)Jae), JAX stock 006965). Unless noted otherwise, mice were housed in the CCHMC Animal Medicine Facility in a clean room set at 22°C and 45% humidity on a daily 12h light/dark cycle, and kept in ventilated racks fed ad libitum with a standard chow diet (5053 -PicoLab® Rodent Diet 20), with bedding changed every two weeks. See figure legend for specific age and number of mice used. All animal experiments were approved by the CCHMC IACUC.

### Tissue dissection

Tissues were carefully dissected to avoid surrounding tissue contamination. Mice were dissected early morning without further alteration, unless noted in the figure legend. Adipose tissue notation and anatomical location (see Figure 1A): dorsal asWAT (dorsal anterior subcutaneous WAT; thin layer covering iBAT dorsally), ventral asWAT (ventral anterior subcutaneous WAT; ventral area of the asWAT), psWAT (posterior subcutaneous WAT; also called inguinal, extends from the dorsal area to the ventral), rWAT (retroperitoneal WAT; located behind the kidneys attached to the dorsal wall of the abdominal cavity), mWAT (mesenteric WAT; a thin, web-like sheet of tissue surrounding the intestines), pgWAT (perigonadal WAT; stems from the testes and epididymis in males), iBAT (interscapular BAT; dorsally facing BAT with pyramidal shape BAT depot), sBAT (subscapular BAT; disperse depot located under the scapula or axillary area), cBAT (cervical BAT; strip of BAT on the deep neck area), aaBAT (aortic arch BAT; small BAT depot on the aortic arch), taBAT (thoracic aorta BAT; lining the descending thoracic aorta), prBAT (perirenal BAT; located in the hilum of the kidney).

### CARLIN labelling

Cas9+/+;rtTA+/+ males and CARLIN+/+ females were timed mated and pregnancies confirmed by vaginal plug detection, with the morning of plug detection designated E0.5. Pregnant dams received three sequential pulses of doxycycline (Dox, 25 µg Dox per gram of body weight), via retro-orbital injection at E6.5, E9.5, and E13.5. Labeled male offspring were used at 8 weeks of age.

### Tissue histology, imaging, and image quantification

Tissue pieces were fixed in 10% formalin. Embedding, sectioning, and Hematoxylin and Eosin (H&E) staining were done by the CCHMC Pathology Core facility. Images were acquired in an Olympus BX51 equipped with a Nikon DS-Fi3 camera. Quantifications of nuclei number and cell size were done using ImageJ.

### RNA-seq analysis

RNA-seq reads were aligned to the UCSC mouse genome 10 mm using the STAR aligner^51^. Only uniquely aligned reads were used for downstream analysis. Raw read counts for each gene were measured using FeatureCounts^52^ with the option “-s 2 -O --fracOverlap 0.8”. Significantly variable genes across all depots were selected using DESeq2^53^ via an LRT test. Genes with FDR < 0.05 and normalized maximum expression > 0.5 RPKM were selected as variable genes. Then, the top 5000 significant genes by FDR were selected for clustering analysis. Eight distinct clusters were identified and were subject to gene ontology analysis using Enrichr^54^.

RNA-seq samples were clustered using the Spearman correlation coefficient as a similarity measure. Genes were clustered using the Pearson correlation coefficient of Z-score samples of log2-normalized expression levels

### CARLIN library preparation and analysis

CARLIN amplicon libraries were generated from bulk adipose tissue RNA, as previously described^25^. Raw paired-end reads were merged using PEAR^55^ and processed through the CARLIN computational pipeline (https://gitlab.com/hormozlab/carlin), which performs custom alignment incorporating a site-dependent cost function, sequence error correction, UMI denoising, and consensus allele calling. Alleles whose observed frequency did not significantly exceed their expected frequency in the reference allele bank were discarded using the Benjamini-Hochberg procedure (FDR = 0.05). Consensus lineage trees were computed from 10,000 stochastic reconstructions, and pairwise tissue distances were calculated as described^25,56^.

### Discordance matrix analysis

To harmonize the gene expression matrix to the CARLIN tissue distance matrix, the CARLIN distances were converted to similarities using linear rescaling. Next, both matrices were z-score normalized using only the off-diagonal elements. For each pair of depots discordance was calculated as Δ = Z_expression − Z_lineage^57^. To assess whether the two matrices are globally correlated, a Mantel test was performed using Spearman rank correlation on the upper triangular elements, with significance assessed by 9,999 permutations^58^.

### CARLIN editing quantification

CARLIN libraries from total RNA from multiple tissues were processed in an Agilent Fragment Analyzer to determine sequence size from multiple tissues. Editing was quantified as a percentage of the edited sequences (size other than the original construct) versus the original size.

### Software used in figures

All graphs were made in GraphPad. Illustrations were made in BioRender. Figures were made in Inkscape. Due to legal requirements, BioRender is acknowledged in the figure legends.

### Quantification and statistical analysis

Data are presented as mean+s.e.m., unless stated otherwise. Statistical analysis used is presented in figure legends. No pre-test was used to choose sample size. The number of mice used per experiment is stated in each figure legend. Statistical analysis was done using GraphPad Prism or R. In all panels, *P < 0.05, **P < 0.01, ***P < 0.001.

### Data availability statement

Datasets generated in this study have been deposited in the NCBI Gene Expression Omnibus database under accession codes GSE348628 and GSE348629.

## Acknowledgements

This work was supported by grants from the NIH (R21OD031907 to JSG, R21OD034470 to JSG, R01DK144589 to JSG and HWL). OI was supported by a JSPS Overseas Research Fellowship (grant number 202360399). This project was supported in part by NIH P30 DK078392 of the Digestive Diseases Research Core Center in Cincinnati. The content is solely the responsibility of the authors and does not necessarily represent the official views of the NIH. We acknowledge the comparative medicine division, the genomics sequencing facility (RRID: SCR_022630) and the integrated pathology research facility (RRID: SCR_022637) at CCHMC. We thank all members of the Sanchez-Gurmaches lab for valuable discussions.

## Authors Contributions

Conceptualization and study design: JSG. Data collection: OI performed most experiments; MG contributed to mouse husbandry and experiments; JSG and AS contributed to morphological and gene expression experiments. Data analysis: OI, CA, MAS, HWL, JSG. Omic data analysis: HWL. Data interpretation: JSG. Supervision: JSG. Manuscript writing – Original draft: JSG. Manuscript writing – Review and editing: all authors. All authors approved the final manuscript.

## Declaration of generative AI and AI-assisted technologies in the writing process

During the preparation of this work the author(s) used Claude.ai in order to improve readability. After using this tool/service, the author(s) reviewed and edited the content as needed and take(s) full responsibility for the content of the publication.

---

**Figure S1:**
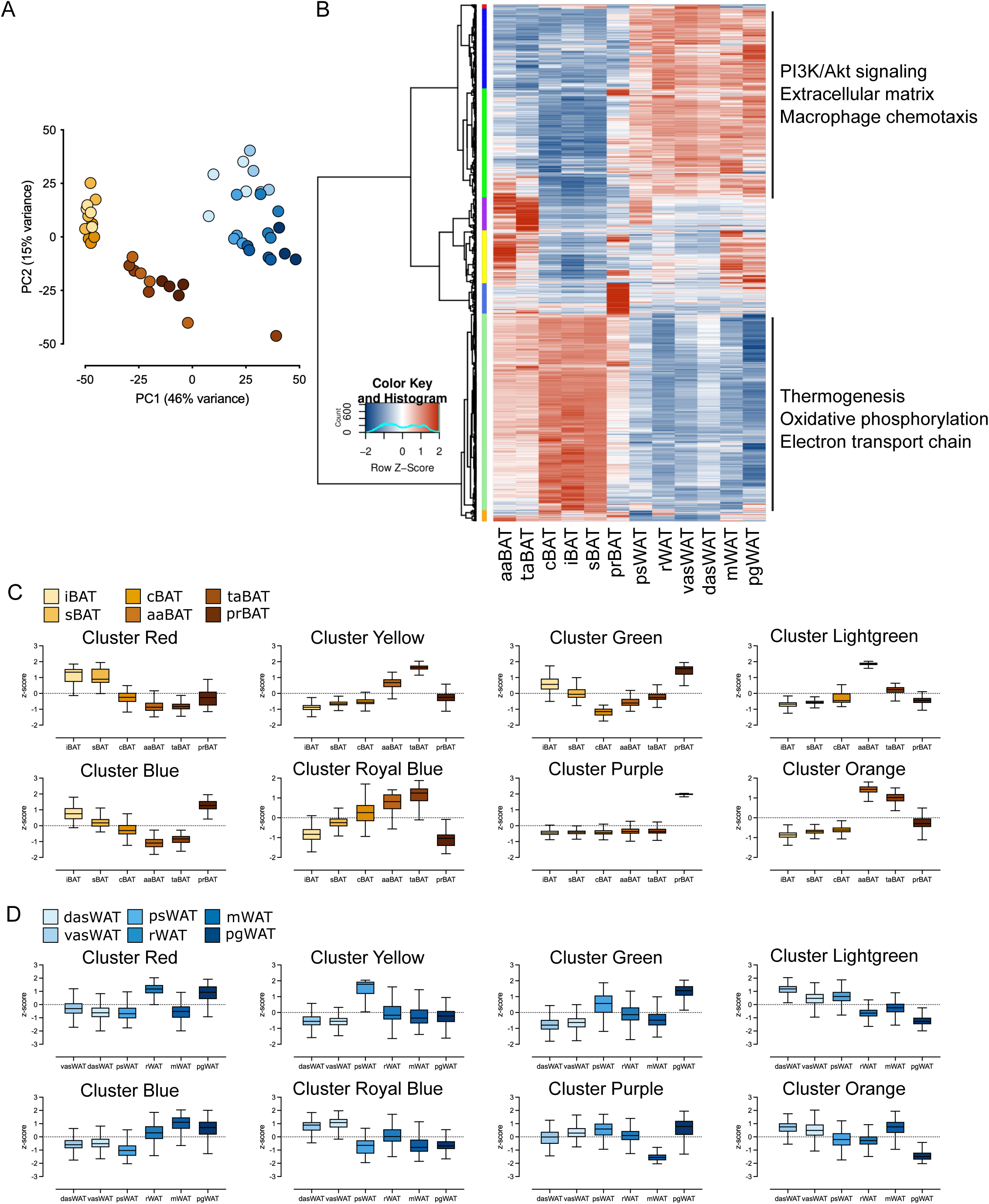
Per-module Z-score distributions across BAT and WAT co-expression clusters. (A) PCA analysis of BAT and WAT depots of 8-weeks-old C57Bl/6J male. Each point represents one biological replicate, colored by depot. n= 4 per depot except pgWAT (n=3). (B) Hierarchical clustering of averaged depot transcriptomes of BAT and WAT depots of 8-weeks-old C57Bl/6J male. Co-expression modules by unsupervised clustering are indicated by the colored bar at left. n= 4 per depot except pgWAT (n=3). (C) Z-score plots between BAT depots among clusters. (D) Z-score plots between WAT depots among clusters.

**Figure S2:**
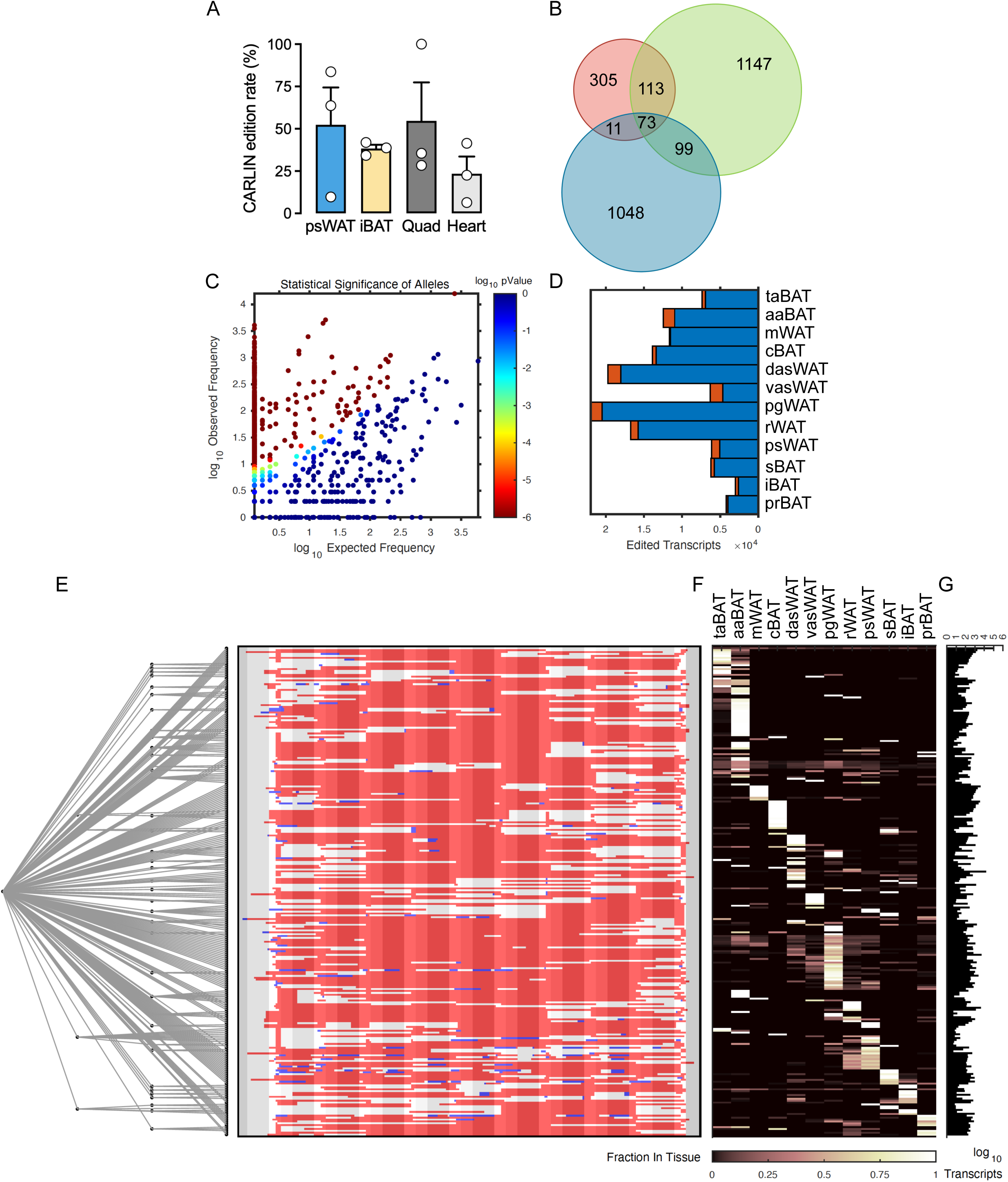
Quality metrics and allele-level analysis of the CARLIN lineage barcoding. (A) CARLIN edition rate in various tissues. Quad: quadriceps. n=3. (B) Overlap of unfiltered barcodes between replicates. (C) A representative scatterplot of observed versus expected allele frequencies. (D) A representative bar graph of the number of edited transcripts found in different adipose tissues after running the CARLIN pipeline (blue and red bars together) and after screening for significant alleles (blue bar) from plot in (B). (E) A representative consensus tree of the sequence of the CARLIN transgene that accounts for 95% of edited transcripts obtained from 10,000 simulations. The ten in tandem target sites in the CARLIN transgene (conserved site (13bp), cutsite (7bp), PAM+linker (7bp)) are represented by a grey mask. Deletions are marked in red. Insertions are marked in blue. (F) A representative distribution of number of transcripts corresponding to each allele across adipose tissues. (G) A representative histogram of total transcript counts across all adipose tissues for each allele.

## References

1 Rosen, E. D. & Spiegelman, B. M. What we talk about when we talk about fat. Cell 156, 20–44 (2014). 10.1016/j.cell.2013.12.012

2 Cinti, S. Adipocyte differentiation and transdifferentiation: plasticity of the adipose organ. Journal of endocrinological investigation 25, 823–835 (2002). 10.1007/BF03344046

3 Sebo, Z. L. & Rodeheffer, M. S. Assembling the adipose organ: adipocyte lineage segregation and adipogenesis in vivo. Development (Cambridge, England) 146 (2019). 10.1242/dev.172098

4 Gesta, S., Tseng, Y. H. & Kahn, C. R. Developmental origin of fat: tracking obesity to its source. Cell 131, 242–256 (2007). 10.1016/j.cell.2007.10.004

5 Lee, K. Y. et al. Developmental and functional heterogeneity of white adipocytes within a single fat depot. Embo j 38 (2019). 10.15252/embj.201899291

6 Young, P., Arch, J. R. & Ashwell, M. Brown adipose tissue in the parametrial fat pad of the mouse. FEBS letters 167, 10–14 (1984).

7 Cousin, B. et al. Occurrence of brown adipocytes in rat white adipose tissue: molecular and morphological characterization. Journal of cell science 103 **( Pt** **4**), 931–942 (1992).

8 Despres, J. P. et al. Regional distribution of body fat, plasma lipoproteins, and cardiovascular disease. *Arteriosclerosis (Dallas*, Tex.) 10, 497–511 (1990).

9 Despres, J. P. Body fat distribution and risk of cardiovascular disease: an update. Circulation 126, 1301–1313 (2012). 10.1161/CIRCULATIONAHA.111.067264

10 Baik, I. et al. Adiposity and mortality in men. American Journal of Epidemiology 152, 264– 271 (2000). 10.1093/aje/152.3.264

11 Pischon, T. et al. General and Abdominal Adiposity and Risk of Death in Europe. New England Journal of Medicine 359, 2105–2120 (2008). 10.1056/NEJMoa0801891

12 Zhang, C., Rexrode, K. M., van Dam, R. M., Li, T. Y. & Hu, F. B. Abdominal obesity and the risk of all-cause, cardiovascular, and cancer mortality -Sixteen years of follow-up in US women. Circulation 117, 1658–1667 (2008). 10.1161/circulationaha.107.739714

13 Pittenger, M. F. et al. Multilineage potential of adult human mesenchymal stem cells. *Science (New York*, N.Y.) 284, 143–147 (1999). 10.1126/science.284.5411.143

14 Wassermann, F. The development of adipose tissue. Comprehensive Physiology 1965, 87–100 (1965). 10.1002/j.2040-4603.1965.tb01226.x

15 Wells, H. G. ADIPOSE TISSUE, A NEGLECTED SUBJECT. Journal of the American Medical Association 114, 2177–2183 (1940). 10.1001/jama.1940.02810220001001

16 Sanchez-Gurmaches, J. & Guertin, D. A. Adipocytes arise from multiple lineages that are heterogeneously and dynamically distributed. Nature communications 5, 4099 (2014). 10.1038/ncomms5099

17 Fu, M. et al. Neural Crest Cells Differentiate Into Brown Adipocytes and Contribute to Periaortic Arch Adipose Tissue Formation. *Arterioscler Thromb Vasc Biol*, Atvbaha119312838 (2019). 10.1161/atvbaha.119.312838

18 Heider, S. et al. The dorsal aortic compartment is a developmental source of brown adipose tissue in mice. Nature communications 17, 286 (2026). 10.1038/s41467-025-68147-9

19 Seale, P. et al. PRDM16 controls a brown fat/skeletal muscle switch. Nature 454, 961– 967 (2008). 10.1038/nature07182

20. Sebo, Z. L., Jeffery, E., Holtrup, B. & Rodeheffer, M. S. A mesodermal fate map for adipose tissue. Development (Cambridge, England) 145 (2018). 10.1242/dev.166801

21 Sanchez-Gurmaches, J. et al. PTEN loss in the Myf5 lineage redistributes body fat and reveals subsets of white adipocytes that arise from Myf5 precursors. Cell metabolism 16, 348–362 (2012). 10.1016/j.cmet.2012.08.003

22 Shan, T. et al. Distinct populations of adipogenic and myogenic Myf5-lineage progenitors in white adipose tissues. Journal of lipid research 54, 2214–2224 (2013). 10.1194/jlr.M038711

23 Wang, W. et al. Ebf2 is a selective marker of brown and beige adipogenic precursor cells. Proc Natl Acad Sci U S A 111, 14466–14471 (2014). 10.1073/pnas.1412685111

24. Billon, N., et al. The generation of adipocytes by the neural crest. Development (Cambridge, England) 134, 2283–2292 (2007). 10.1242/dev.002642

25 Bowling, S. et al. An Engineered CRISPR-Cas9 Mouse Line for Simultaneous Readout of Lineage Histories and Gene Expression Profiles in Single Cells. Cell 181, 1410– 1422.e1427 (2020). 10.1016/j.cell.2020.04.048

26 de Jong, J. M., Larsson, O., Cannon, B. & Nedergaard, J. A stringent validation of mouse adipose tissue identity markers. American journal of physiology. Endocrinology and metabolism 308, E1085–1105 (2015). 10.1152/ajpendo.00023.2015

27 Sanchez-Gurmaches, J., Hsiao, W. Y. & Guertin, D. A. Highly selective in vivo labeling of subcutaneous white adipocyte precursors with Prx1-Cre. Stem cell reports 4, 541–550 (2015). 10.1016/j.stemcr.2015.02.008

28 Cannon, B. & Nedergaard, J. Brown adipose tissue: function and physiological significance. Physiological reviews 84, 277–359 (2004). 10.1152/physrev.00015.2003

29 Cinti, S. The adipose organ. Prostaglandins, leukotrienes, and essential fatty acids 73, 9– 15 (2005). 10.1016/j.plefa.2005.04.010

30 Tseng, Y. H., Cypess, A. M. & Kahn, C. R. Cellular bioenergetics as a target for obesity therapy. Nat Rev Drug Discov 9, 465–482 (2010). 10.1038/nrd3138

31 Goodwin, L. O. et al. Large-scale discovery of mouse transgenic integration sites reveals frequent structural variation and insertional mutagenesis. Genome research 29, 494–505 (2019). 10.1101/gr.233866.117

32 Nicholls, P. K., Bellott, D. W., Cho, T. J., Pyntikova, T. & Page, D. C. Locating and Characterizing a Transgene Integration Site by Nanopore Sequencing. G3 (Bethesda) 9, 1481–1486 (2019). 10.1534/g3.119.300582

33 Cain-Hom, C. et al. Efficient mapping of transgene integration sites and local structural changes in Cre transgenic mice using targeted locus amplification. Nucleic acids research 45, e62 (2017). 10.1093/nar/gkw1329

34 Dinsmore, C. J., Ke, C. Y. & Soriano, P. The Wnt1-Cre2 transgene is active in the male germline. Genesis (New York, N.Y. : 2000) 60, e23468 (2022). 10.1002/dvg.23468

35 Liu, S., Lockhart, J. R., Fontenard, S., Berlett, M. & Ryan, T. M. Mapping the Chromosomal Insertion Site of the GFP Transgene of UBC-GFP Mice to the MHC Locus. J Immunol 204, 1982–1987 (2020). 10.4049/jimmunol.1901338

36 Lee, S., Kuramochi-Miyagawa, S., Nagamori, I. & Nakano, T. Effects of transgene insertion loci and copy number on Dnmt3L gene silencing through antisense transgene-derived PIWI-interacting RNAs. Rna 28, 683–696 (2022). 10.1261/rna.078905.121

37 Laboulaye, M. A., Duan, X., Qiao, M., Whitney, I. E. & Sanes, J. R. Mapping Transgene Insertion Sites Reveals Complex Interactions Between Mouse Transgenes and Neighboring Endogenous Genes. Front Mol Neurosci 11, 385 (2018). 10.3389/fnmol.2018.00385

38 Lewis, A. E., Vasudevan, H. N., O’Neill, A. K., Soriano, P. & Bush, J. O. The widely used Wnt1-Cre transgene causes developmental phenotypes by ectopic activation of Wnt signaling. Developmental biology 379, 229–234 (2013). 10.1016/j.ydbio.2013.04.026

39 Perl, A. J. et al. Reduced Nephron Endowment in Six2-TGCtg Mice Is Due to Six3 Misexpression by Aberrant Enhancer-Promoter Interactions in the Transgene. J Am Soc Nephrol 35, 566–577 (2024). 10.1681/asn.0000000000000324

40 Zheng, H. et al. A Cre-driver rat model for anatomical and functional analysis of glucagon (Gcg)-expressing cells in the brain and periphery. Molecular metabolism 66, 101631 (2022). 10.1016/j.molmet.2022.101631

41 Yong, C. S. et al. Embryonic Lethality in Homozygous Human Her-2 Transgenic Mice Due to Disruption of the Pds5b Gene. PloS one 10, e0136817 (2015). 10.1371/journal.pone.0136817

42 Álvarez-Aznar, A. et al. Tamoxifen-independent recombination of reporter genes limits lineage tracing and mosaic analysis using CreER(T2) lines. Transgenic research 29, 53– 68 (2020). 10.1007/s11248-019-00177-8

43 Reinert, R. B. et al. Tamoxifen-Induced Cre-loxP Recombination Is Prolonged in Pancreatic Islets of Adult Mice. PloS one 7, e33529 (2012). 10.1371/journal.pone.0033529

44 Ye, R. et al. Impact of tamoxifen on adipocyte lineage tracing: Inducer of adipogenesis and prolonged nuclear translocation of Cre recombinase. Molecular metabolism 4, 771– 778 (2015). 10.1016/j.molmet.2015.08.004

45 Angueira, A. R. et al. Defining the lineage of thermogenic perivascular adipose tissue. Nat Metab 3, 469–484 (2021). 10.1038/s42255-021-00380-0

46 Wang, Y. et al. The role of somatosensory innervation of adipose tissues. Nature 609, 569–574 (2022). 10.1038/s41586-022-05137-7

47 Chi, J. et al. Three-Dimensional Adipose Tissue Imaging Reveals Regional Variation in Beige Fat Biogenesis and PRDM16-Dependent Sympathetic Neurite Density. Cell metabolism 27, 226–236 e223 (2018). 10.1016/j.cmet.2017.12.011

48 Willows, J. W., Blaszkiewicz, M. & Townsend, K. L. The Sympathetic Innervation of Adipose Tissues: Regulation, Functions, and Plasticity. Comprehensive Physiology 13, 4985–5021 (2023). 10.1002/cphy.c220030

49 Escobedo, N. & Oliver, G. The Lymphatic Vasculature: Its Role in Adipose Metabolism and Obesity. Cell metabolism 26, 598–609 (2017). 10.1016/j.cmet.2017.07.020

50 Jespersen, N. Z. et al. Heterogeneity in the perirenal region of humans suggests presence of dormant brown adipose tissue that contains brown fat precursor cells. Molecular metabolism 24, 30–43 (2019). 10.1016/j.molmet.2019.03.005

51 Dobin, A. et al. STAR: ultrafast universal RNA-seq aligner. *Bioinformatics (Oxford*, England) 29, 15–21 (2013). 10.1093/bioinformatics/bts635

52 Liao, Y., Smyth, G. K. & Shi, W. featureCounts: an efficient general purpose program for assigning sequence reads to genomic features. *Bioinformatics (Oxford*, England) 30, 923– 930 (2014). 10.1093/bioinformatics/btt656

53 Love, M. I., Huber, W. & Anders, S. Moderated estimation of fold change and dispersion for RNA-seq data with DESeq2. Genome Biol 15, 550 (2014). 10.1186/s13059-014-0550-8

54 Kuleshov, M. V. et al. Enrichr: a comprehensive gene set enrichment analysis web server 2016 update. Nucleic acids research 44, W90–97 (2016). 10.1093/nar/gkw377

55 Zhang, J., Kobert, K., Flouri, T. & Stamatakis, A. PEAR: a fast and accurate Illumina Paired-End reAd mergeR. *Bioinformatics (Oxford*, England) 30, 614–620 (2014). 10.1093/bioinformatics/btt593

56 Feng, J. et al. ESTIMATION OF CELL LINEAGE TREES BY MAXIMUM-LIKELIHOOD PHYLOGENETICS. Ann Appl Stat 15, 343–362 (2021). 10.1214/20-aoas1400

57 Legendre, P. Comparison of permutation methods for the partial correlation and partial mantel tests. Journal of Statistical Computation and Simulation 67, 37–73 (2000). 10.1080/00949650008812035

58 Mantel, N. The detection of disease clustering and a generalized regression approach. Cancer research 27, 209–220 (1967).

